# Consideration of a Liquid mutation-accumulation Experiment to Measure Mutation Rates by Successive Serial Dilution

**DOI:** 10.1101/2023.08.31.555790

**Authors:** Stephan Baehr, Wei-Chin Ho, Sam Perez, Alyssa Cenzano, Katelyn Hancock, Lea Patrick, Adalyn Brown, Sam Miller, Michael Lynch

## Abstract

The mutation-accumulation (MA) experiment is a fixture of evolutionary biology, though it is laborious to perform. MA experiments typically take between months and years to acquire sufficient mutations to measure DNA mutation rates and mutation spectra. MA experiments for many organisms rely on colony formation on agar plates and repetitive streaking, an environment which at first glance appears somewhat contrived, a poor imitation of real environmental living conditions. We propose that a fully liquid-phase mutation-accumulation experiment may at times more accurately reflect the environment of an organism. We note also that whereas automation of streaking plates is a daunting prospect, automation of liquid handling and serial dilution is already commonplace. In principle, this type of MA experiment can be automated so as to reduce the human capital requirements of measuring mutation rates. We demonstrate that a liquid MA recapitulates the mutation rate estimated for MMR- *E. coli* in liquid LB culture vs. plate LB culture. We detect a modified mutation spectrum with a transition skew of 4:1 of A:T→G:C vs G:C→A:T mutations, highlighting the potential role of tautomerization as a DNA mutation mechanism. We also find that using a plate reader to measure OD600 as a proxy for cell growth to be incapable of measuring carrying capacity for MA lines burdened with many mutations.

## 1 Introduction

Mutation rates describe the input of heritable variation into an organism. Because the average mutation is deleterious[3], evolution pushes DNA mutation rates to the lower limits of natural selection[14]. DNA mutation events occur between 1 in 100,000,000 bases to 1 in 1,000,000,000,000 bases per cell division[14], making their identification within sequenced genomes equivalent to the search for a needle in a rather large haystack. Techniques that amplify the signal of mutation events allow a general means of detection; the fluctuation test, first demonstrated in 1943[13, 4], uses a selective agent to make certain mutation events stand out, from which mutation rates can be estimated [11, 4, 29]. With the advent of high-throughput sequencing, the mutation-accumulation (MA) experiment[15, 14], which employs recurrent single-cell bottlenecks typically on some surface like an agar plate, has become the gold standard for the empirical estimation of mutation. The analysis of a MA is straightforward, requiring the counting of new mutations arisen over a period of single-cell bottlenecks, and an estimation of the number of cell divisions, so as to come up with the numerator and a denominator that comprises a mutation rate.

Some criticism has been directed to MA experiments. An agar-based or other petri dish likely bears little resemblance to most cellular growth environments, from *E. coli* [7] to human cells. Oxygen is not a constant presence in many organism environments, though it is almost universally present in MA experiments unless expressly engineered out[20]. Most cells used for MA experiments grow into colonies of some sort, a growth structure that is nutrient poor and perhaps different from an organism’s natural growth structure, for example in a human gut where we might expect some level of nutrient flow or churn. Starvation has been proposed as a state that may be mutagenic[23, 5], though typically the time frames are longer than 24 hours, and the experiments have trouble resolving the difference between higher mutation rates or more cell divisions occurring[8, 28]. Depending on how a researcher picks and spreads their colonies, edge effects may skew cell division estimates, as some parts of a colony experience more divisions than others[26]. Though single-cell bottlenecks are expected to overwhelm any but the most stringent selection (lethality), the harmonic mean effective population size is closer to 14 than 1[25, 16]; Several authors propose selection is a significant factor in a generic MA experiment, either throughout or in at least the first two weeks[2]. Colony choice instigated by researchers during serial bottlenecking is also at risk of researcher bias, in that humans tend to pick the colonies they can see, rather than just any colony. Humans can also select, consciously or without realizing, a certain colony phenotype, perhaps skewing the distribution of mutations observed in an experiment. The sum of these criticisms is not entirely without merit; there is certainly some selection, positive and purifying, occurring in a MA experiment[9, 22]. Regardless, it is difficult to expect mutation-rate estimations are off by more than a factor of 2; for this to be the case, half of all mutations would have to be lethal.

Some of the above concerns of bias may be addressed by a MA experiment with a less structured environment. Researchers have long appreciated the ability of serial dilution to reach countable sub-samples of bacterial populations. In principle, some level of dilution can be achieved so that half of all wells inoculated by a dilution contain a viable cell, and half do not. By reaching this level of dilution or lower, relatively high confidence can be maintained that a single viable bacterium was the source of the growth observed in a well 24 hours later. Moreover, wild-type *E. coli* can reach carrying capacity within 12-14 hours, given a cell division rate of 20-30 minutes[24]. This suggests that a fixed fitness burden of up to 50 percent may reasonably be tolerated without affecting our ability to select it. If carrying capacity remains relatively constant, the number of cell divisions should be quite easy to determine relative to a plate MA, which typically develops wildly different colony sizes over time[12]. The liquid-based single-cell bottleneck eschews the semi-structured agar environment, and reduces the potential risk of artificial selection imposed by researchers. It is also readily recognizeable that a liquid MA lends itself to automation by liquid handling robots, in contrast to the agar-based MA’s that escape easy conversion to robotic control. If this automation could be achieved, one of the most laborious aspects of obtaining mutation rates by an MA experiment could be made more tolerable. We did not automate our liquid MA, but the point remains enticing for future work.

To compare and contrast the output of MA experiments run by liquid serial dilution or by standard agar plates, we chose to use a well-studied *E. coli* strain, MutL- or MMR-E coli, K12, MG1655, originating from the lab of Pat Foster, a strain which has already been characterized by a standard MA experiment, twice[12, 28]. This strain is particularly useful because it features a distinct mutation spectrum, as well as an elevated mutation rate, at least 100-fold higher than WT. The elevated mutation rate allows for a temporally shorter MA with relatively few samples to yield an abundance of mutations, along with measureable phenotypes. We ran a plate and a liquid MA in tandem from the same starting single colony for 20 days, and afterward sequenced the DNA. We find a modest difference in the mutation spectrum of *E. coli*, but moreso are struck by the invariance in mutation-rate estimated from the experiment between growth types.

## 2 Methods

Frozen stocks of *E. coli* MMR- were thawed and streaked upon an LB plate (LB Agar, Miller). 24 hours later, a single colony was chosen, from which 1 10 mL liquid LB tube (LB Broth, Miller) was inoculated while 4 additional LB plates were streaked. From the four agar plates, 40 colonies were chosen to be the set of 40 MA lines, and comprised the first day of the plate MA. For each of these 40 lines, per day, every 24 hours, a single colony was selected and streaked on half of a LB plate, with two or three streaks to obtain single colonies. The first streak was a toothpick line (autoclaved toothpicks), and one or two consecutive streaks thereafter came from autoclaved wooden sticks. The last colony of a streak was always chosen for transfer, unless forced by poor streaking to pick the last convincing single colony. Plates were loaded into a 37°C incubator and grown overnight for 24 hours, +/-2 hours, before streaking was repeated on new LB agar plates the next day.

From the 10 mL liquid LB tube, first an ancestor line frozen stock was made and then 16 wells of 20 uL were taken and run through serial dilution. Briefly, *E. coli* were serially diluted to the -5 and -6 wells by standard methods and multichannel pipettors in 180 uL 1x PBS in 96 well plates. Then, 100 uL of the -5 was added to 900 uL of 1x PBS to create a ”-7” dilution of 1 mL per sample, and the same was done to the -6 well to create a ”-8” dilution of 1 mL per sample. From the -7 dilution, 20 uL was inoculated to a single row of a 96-deepwell plate holding 950 uL of liquid LB, 8 wells. From the -8 dilution, a second row for the same sample was loaded with 50uL, and for a 3rd row the -8 dilution was again used to inoculate 20uL. This results in an order of magnitude scale of dilution in the 96 deepwell plate, where each liquid sample comprises now 24 replicates of varying serial dilution. On a 96-deepwell plate four samples were arrayed, for a total of 4 96-deepwell plates, or 16 replicates, per transfer day. After 24 hours, some of the serial dilution wells have turbid growth, while others remain clear. At the lower serial dilutions, a majority of the wells should be clear. As a rule, the turbid well from the lowest serial dilution would be chosen for the next round of serial dilution. In the event of a tie, the one on the left-hand side was chosen. After 15 days, frozen stocks were made of both the liquid and the plate MA. After noticing an apparent decrease in carrying capacity in one of the liquid MA lines, the experiment was stopped at 20 days for characterization and sequencing. Based on the previous MA experiment[12], we expected 20 days would result in a sufficient number of mutations to analyze the data.

After the MA’s were complete and frozen stocks obtained, DNA was extracted by first growing the frozen stocks overnight in 10 mL tubes and then running the Promega Wizard DNA Extraction Kit, with minor modifications immaterial to results save by increasing yield to greater than 1 ug per sample. Genomic DNA was quantified, and submitted to the Beijing Genomics Institute for their in house library prep and DNB-sequencing.

Raw sequence reads were analyzed by fastqc to confirm a successful sequencing run. Reads were then filtered with Trimmomatic to remove adapter sequences. Reads were then aligned to the *E. coli* K12 reference genome (https://www.ncbi.nlm.nih.gov/nuccore/U00096.2). After alignment and conversion to bam files by SamTools, the mutation caller GATK2 generated VCF files of novel mutations when comparing the ancestor to the evolved lines. Output VCF files were additionally annotated by SnpEff to determine Dn/Ds and Genic/Intergenic ratios.

To estimate the number of cell divisions that occurred over the course of the 20-day experiment, the liquid and plate MAs were handled differently. For the plate MA, 20-day frozen stocks and the ancestor were thawed and grown in LB overnight in 10 mL tubes. This liquid was then streaked on LB plates. After 24 hours, the colonies were resuspended in LB and serially diluted to the -6 plate, and plated to count colony forming units (CFU’s) per mL. An average of the entire set of 16 plate MA samples was used to calculate mean number of cell divisions per day at day 20, the ancestor calculated mean number of cell divisions per day at day 1. The average between the day 1 and mean of day 20 samples was used as an estimate for number of cell divisions per line.

To estimate cell divisions from the liquid MA, 20-day frozen stocks and the ancestor were thawed and grown overnight in LB in 10 mL tubes. After 24 hours, 20 uL aliquodt’s were taken from the overnight cultures and serially diluted down to single cell bottlenecks, before being grown up in 96-deepwell plates, in a method identical to the MA serial dilution. After an additional 24 hours, 4 turbid replicates were chosen for serial dilution to count CFUs. From this we calculated the number of cell divisions per day, at day 1 and day 20. The mean of day 20 cell division estimates, and the mean between the day 1 number of cell divisions and day 20 was used as an estimate for average total number of cell divisions during the experiment.

To phenotype the lines for fitness and carrying capacity using a plate reader, a 96-well plate was used. Two lines and an ancestor were thawed out per day and grown overnight. 100 uL of overnight culture was then added to 10 mL of LB in 15 mL tubes. These 15 mL tubes were inverted 15 times to mix. The mixtures were then loaded onto a 96-well plate, 200 uL per plate, staggered such that rows 1, 4, 7, and 10 received the ancestor, 2, 5, 8, 11 received MA line 1, and 3, 6, 9, 12 received MA Line 2. the 96-well plate was double-sealed with parafilm and loaded into a plate reader for 24 hours of optical-density analysis, at 37°C, with readings taken at a wavelength of 600nm (OD600). Growth rate was obtained by calculating the largest slope of growth between 2 and 10 hours, when averaged across 5 time points. Carrying capacity was calculated at average optical-density at 12 hours, averaged over 5 time points. Data were normalized per day in comparison to the ancestor, and then the relative fitness and relative ”carrying capacity” were graphed in R.

To phenotype the lines for carrying capacity after 12 hours by CFU, all lines were thawed in 10 mL tubes of LB overnight. The lines were then diluted in quadruplicate to 1/100th concentration in a 96-well plate, by adding 2 uL of mixed overnight culture to 198 uL of fresh LB. These plates were then sealed with parafilm in 2 layers and loaded into a 37 degree incushaker shaking at 200RPM, secured by magnets, for 12 hours. After 12 hours, the E coli were serial diluted to the -7 plate and CFUs were obtained by spreading 100 uL on LB plates. Data were processed by collecting CFU estimates, running t tests, and graphing the different CFU averages with standard error of the mean in R.

## 3 Results

The tandem mutation-accumulation experiments in liquid and upon plates of otherwise identical nutrient content are quite nearly identical in estimation of mutation rate per site per generation, and in general agreement with previous mutation-accumulation experiments (Figure 2). This resulted in similar mutation burdens over the course of the experiment (Table 1), with the difference in burden being explained by the liquid MA experiencing slightly more cell divisions per day. Specifically, the liquid MA lines had a higher mutation burden in base pair substitutions (BPS) by the end of the experiment, almost 10%, which can be explained by the cells having experienced 10% more cell divisions (29.9 vs. 26.9 mean cell divisions per line per day).

**Table 1:** Summary Statistics for the Liquid and Plate MA experiments.

| Treatment | Sample Size | Mean BPS per line | Mean generations per transfer | Mutation Rate/site/Gen $\times 10^{-8}$ |
| --- | --- | --- | --- | --- |
| Liquid | 16 | 76.8 (+/- 6.32) | 29.9 | 2.80 (+/- 0.24) |
| Plate | 16 | 71.9 (+/- 6.20) | 26.9 | 2.91 (+/- 0.25) |

**Table 2:**
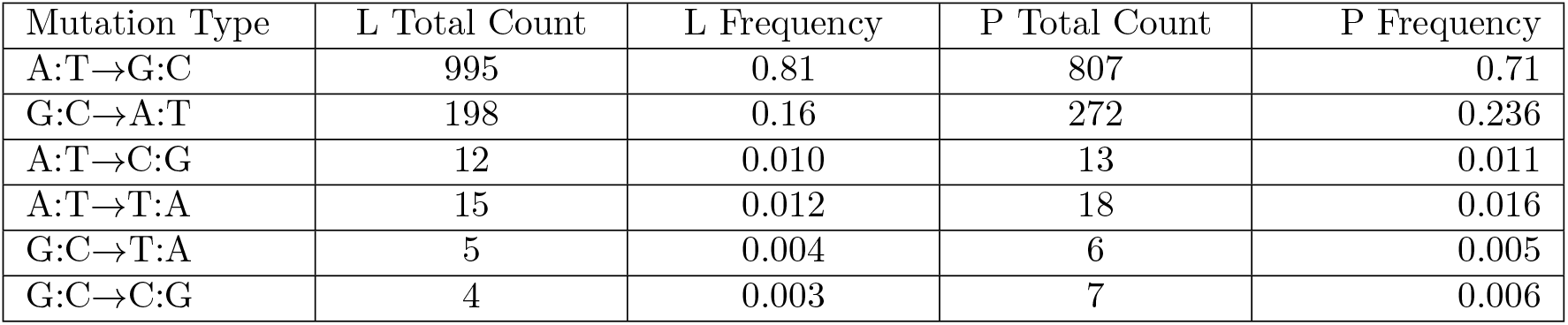
Supplementary Table 1: Raw Data of the Mutation Spectrum.

| Mutation Type | L Total Count | L Frequency | P Total Count | P Frequency |
| --- | --- | --- | --- | --- |
| A:T→G:C | 995 | 0.81 | 807 | 0.71 |
| G:C→A:T | 198 | 0.16 | 272 | 0.236 |
| A:T→C:G | 12 | 0.010 | 13 | 0.011 |
| A:T→T:A | 15 | 0.012 | 18 | 0.016 |
| G:C→T:A | 5 | 0.004 | 6 | 0.005 |
| G:C→C:G | 4 | 0.003 | 7 | 0.006 |

**Figure 1:**
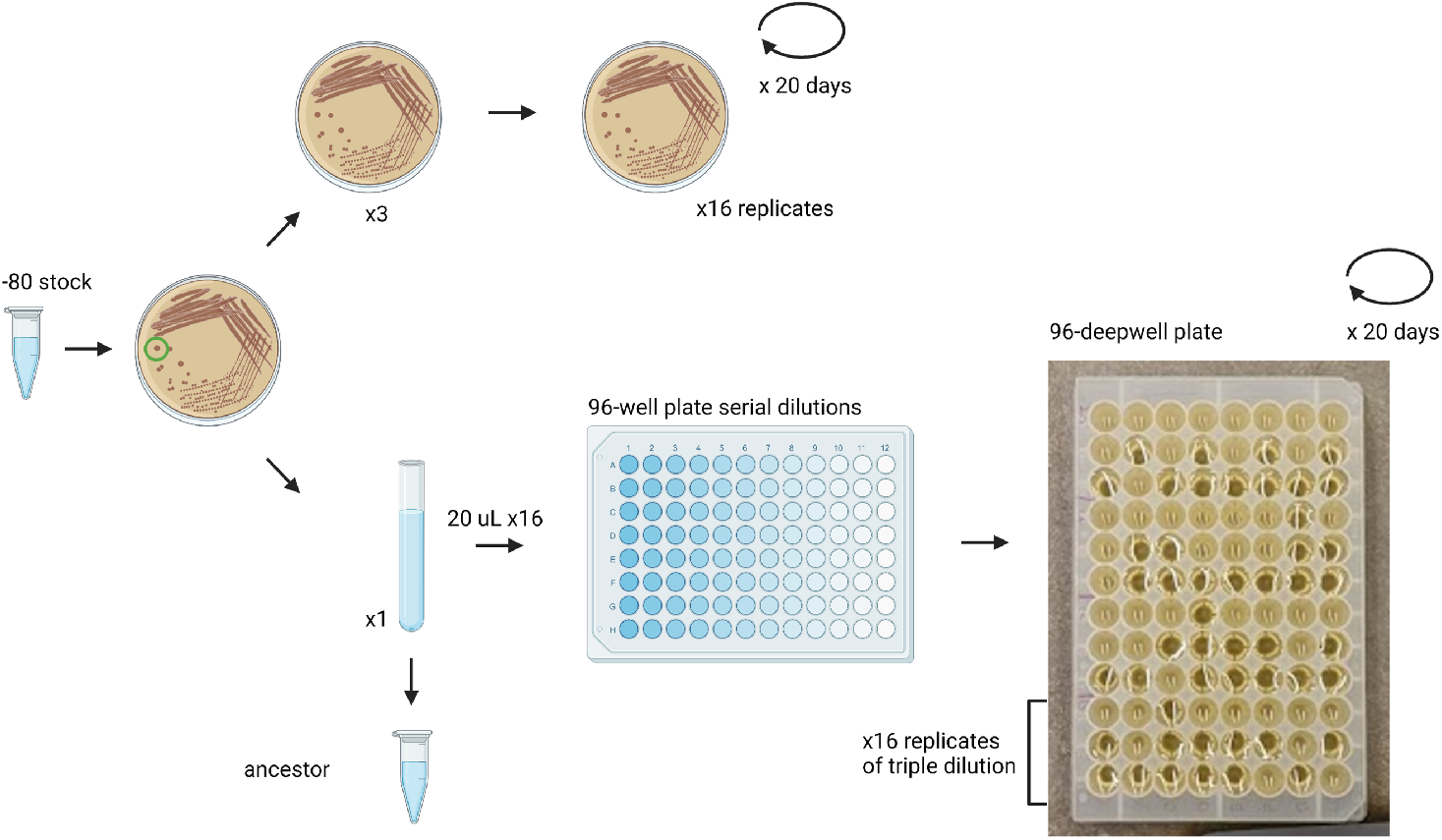
Experimental Design. From a single colony streaked on a plate, liquid and plate MA’s were started in tandem. The plate MA used standard methods. The Liquid MA begins with a serial dilution to the point of getting 0 or 1 cells in a well. Three different serial dilutions are used to ensure sufficient dilution.

**Figure 2:**
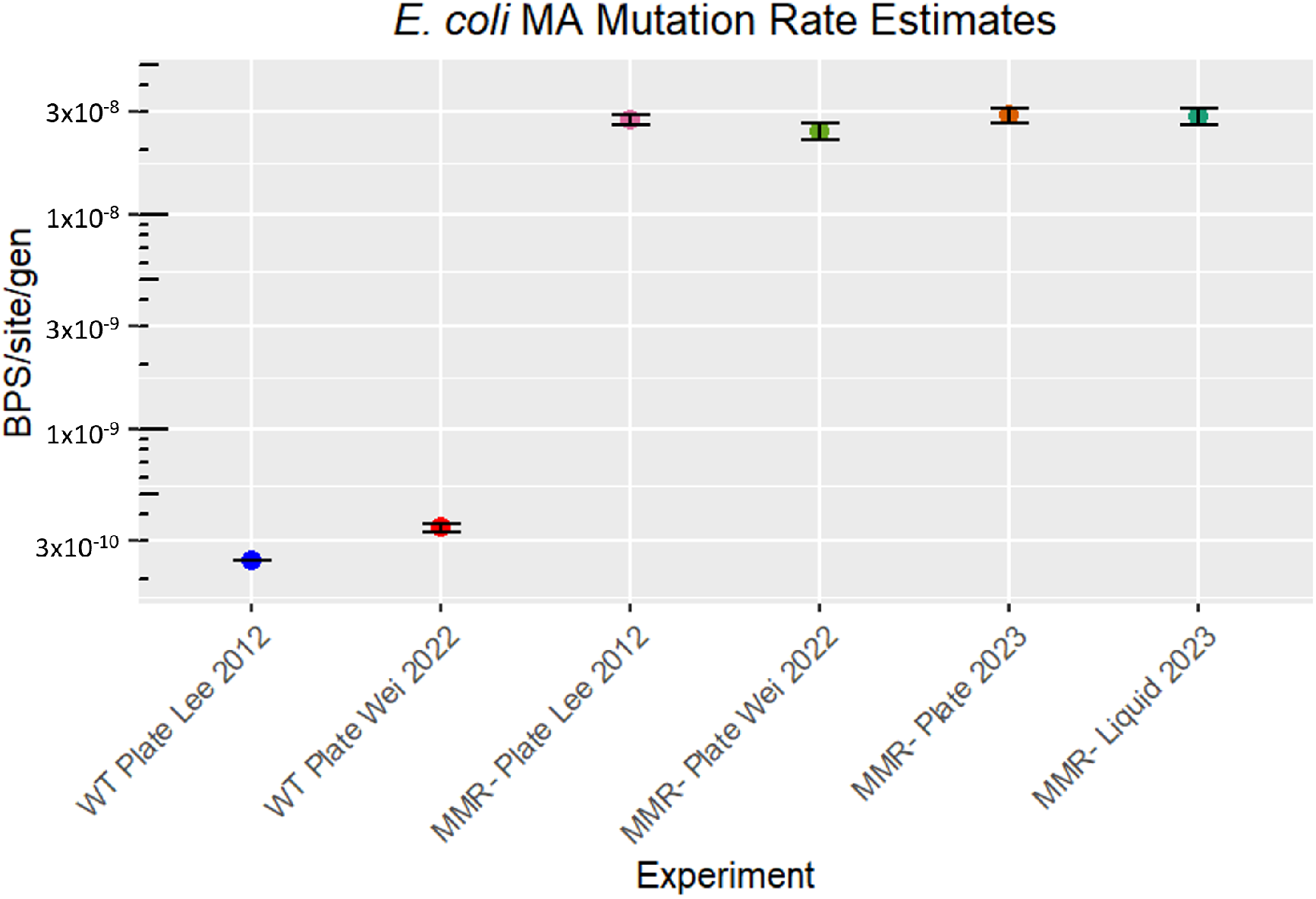
Mutation Rate Estimation. Multiple estimates of the *E. coli* MMR- mutation rate are consistent over time. The liquid and plate experiments produced mutation rate estimates more similar to each other than the difference between plate MA experiments over time.

The mutation spectra are similar to previous estimates from 2012 and 2022 (Figure 3a), although a *χ*-square test detected a significant difference between liquid and plate spectra (Figure 3b) (p = 8.8 × e-8), driven by both a reduction in G:C→ A:T transitions and an increase in A:T →G:C transitions. Though the difference by ratio is modest in appearance, the net effect over 20 days yields a difference of 74 G:C →A:T mutations across summed across the 16 lines (Supplementary Table 1), a 37% increase in occurrence in the plate vs. liquid MA experiments.

**Figure 3:**
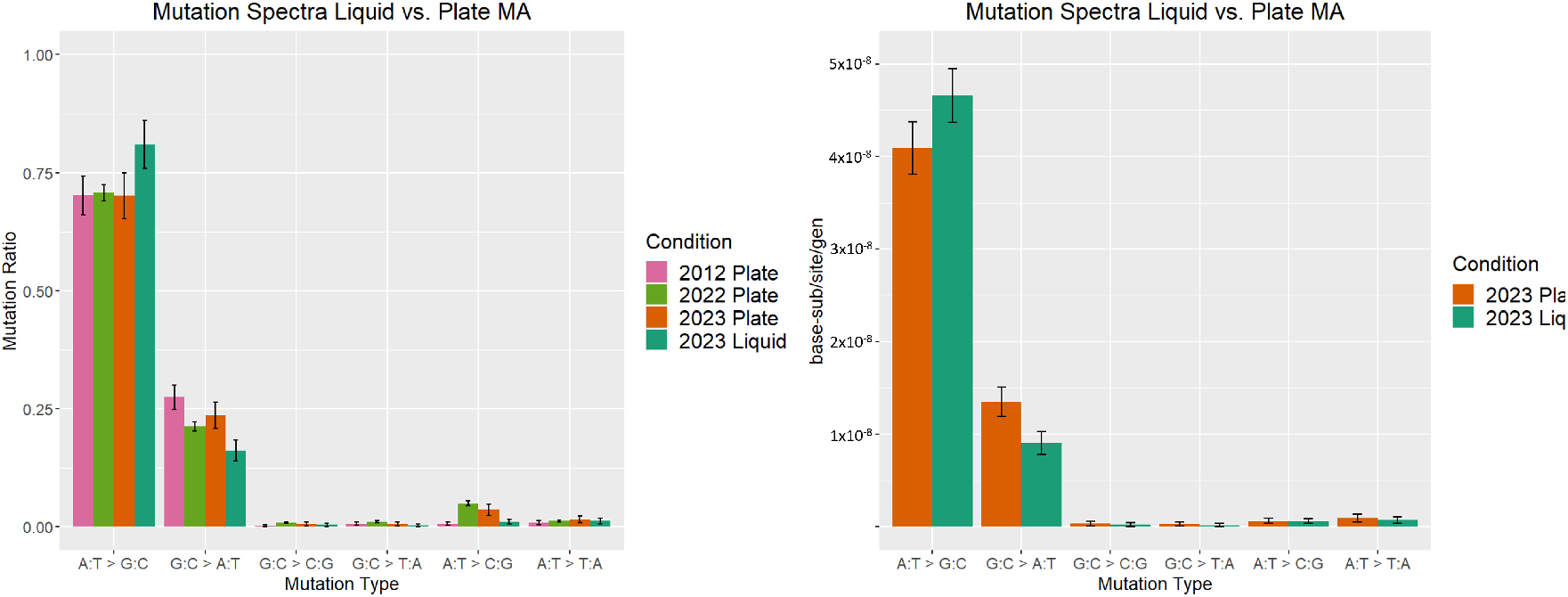
Mutation Spectra of MMR- *E. coli*. A. Across MA experiments the mutation spectrum ratios of MMR- *E. coli* mutations holds relatively constant, particularly in the case of the plate MA. B. Mutation rates rather than mutation ratios. In this graph, the mutations from hypermutator line P35 are omitted.

The liquid MA experienced roughly 3 additional mutations per day (29.9 vs 26.9, p value = 7.662e-11, t-test), a consequence of the 1 mL growth volume allotted per well in the 96-deepwell plate. We note that a difference in the number of cell divisions is not only in count per day but also in variation per colony, which is readily discerned in a comparison of the standard error of the mean in the estimates of number of cells per day (Figure 4). The increased variance, readily discernable by eye in Figure 4a, error bars liquid samples vs. plate samples, can be compared by calculating the coefficient of variation (CV), which normalizes the standard deviation of a sample by its mean value. The difference in CV is statistically significant (Figure 4b, (p value = 0.00014, t-test), and is readily explained by colony dynamics. When colonies are close to one another they experience nutrient competition and grow less quickly. Random differences in spacing between colonies leads to variation in colony size, and thus increased variation in the total number of cell divisions experienced in a day relative to liquid cultures. It is also true that colonies shrink on average over the course over a mutation-accumulation experiment, as mutations accumulate and fitness degrades. This shrinking colony size is readily seen in measurements of fitness, for example in plate line 25 (Figure 4a).

**Figure 4:**
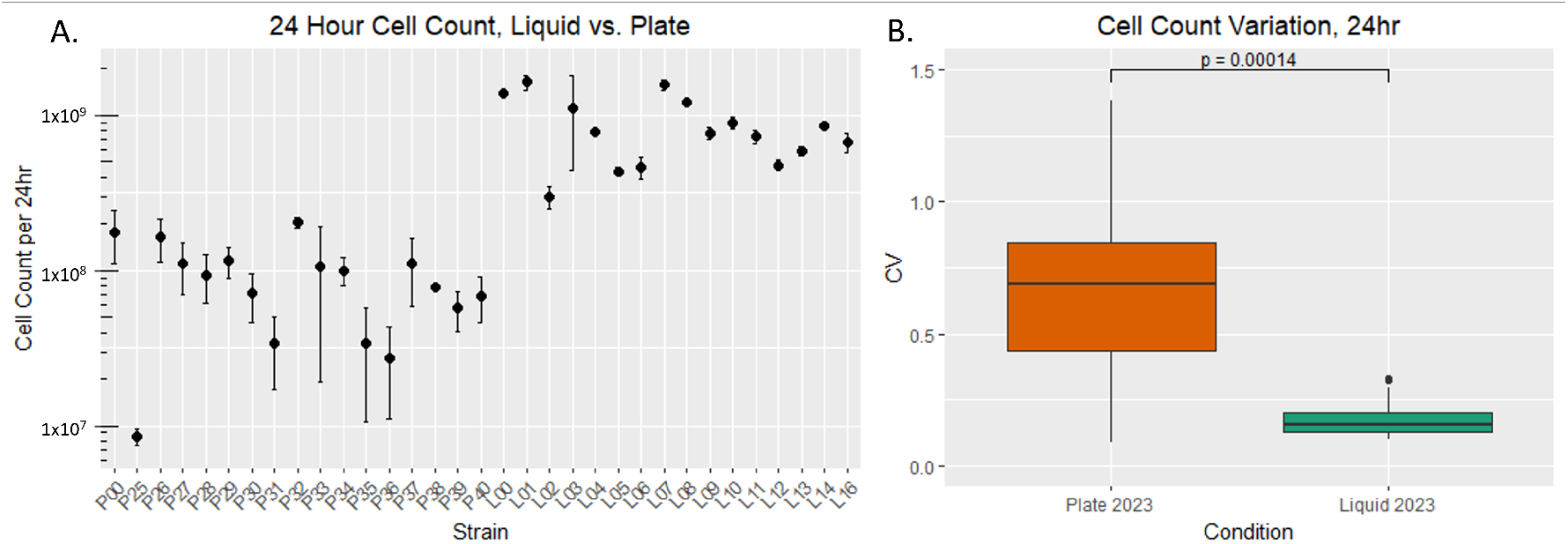
Cell Counts per Day, MMR- *E. coli*. A. CFU estimate of the number of cells present in a culture after 24 hours of growth, from resuspended colonies or from 1 mL liquid cultures for the plate and liquid MA’s, respectively. B. Coefficient of Variation summed over the two MA’s, indicating that there is a greater CV and distribution of CV in the plate MA than the liquid MA.

One of the hallmarks of a successful MA experiment is a decline in mean fitness of the lines[9]. To ensure our MA ran as anticipated, we measured fitness through use of a 96-well plate reader, which determines changes in optical-density over time. In particular, the maximum slope of optical-density change in the first 8 hours of the experiment is sufficient to provide a useful proxy for maximum cell growth rate (Figure 5a, 5b), as has been employed elsewhere. We noticed through the experiments that the MA lines tended to have higher optical-density than the ancestor by 10 hours or so, and we also noticed that the OD readings peaked around 12 hours and seemed to recede some 20% by the end of our growth experiment (5a). Further, we noted that overall, the MA lines had higher OD readings toward the tail of the experiment, relative to the ancestor (Figure 5c). We wondered if the MA lines, in particular the liquid MA lines, had evolved higher carrying capacity by some form of selection. To verify the results from the 96-well plate reader, we ran CFU’s of the 12-hour time-point (5d). We found that despite the optical-density being statistically significantly higher in the MA lines vs. the ancestor, this OD600 absorbance was not due to increased numbers of cells. Thus, we conclude that the OD600 measurement by plate reader of carrying capacity is of limited use, at least in MA lines of hypermutator *E. coli*. Based on the growth phase and CFU counts (Figure 5b, 5d) we calculate the average fitness cost per mutation to be 0.001.

**Figure 5:**
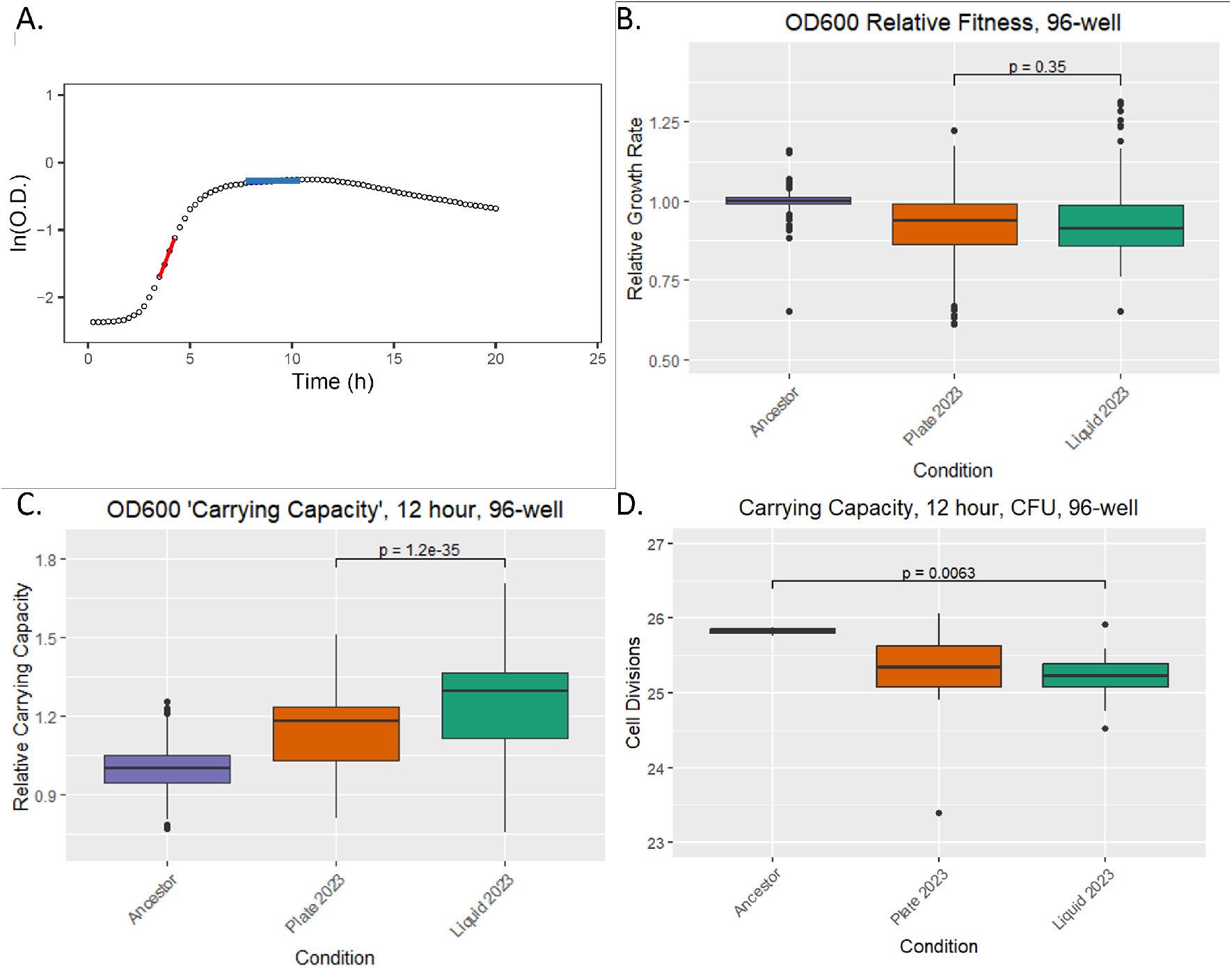
96-well vs. CFU Fitness, Carrying Capacity vs Ancestor in MMR- *E. coli*. A.Example growth curve from a plate reader in a single well, in which culture turbidity is measured at 15min. The red line indicates the calculation of fitness, the blue line demonstrates how carrying capacity is calculated. B. Relative fitness, normalized to ancestral fitness. C. Relative Carrying Capacity at 12hr, normalized to ancestral Carrying Capacity. D. Cell count at 12hr as estimated by CFU.

Through the course of an MA, a distribution of fitness decline is expected and was observed. On day 15 of the liquid MA experiment, one particular line, Liquid line 2, was perceived to have significantly decreased in 24-hour cell count on the day of transfer, moreso than all others (Figure 6a). Time-points of day 14, 15, and 16 were collected for the line, in the hopes of determining which mutation was most responsible for the loss in cell count. Given the mutation rate, number of cell divisions, and genome size, we expect in the neighborhood of 4 mutations to accumulate on average per line per day. Four new mutations were detected between days 14 and 15 (Figure 6b), a result of the bottlenecking process. Two of the mutations are synonymous, and one is intergenic in the middle of an operon, and thus seem unlikely to be responsible for such dramatic phenotypic decline. The 4th mutation, in the CadA gene, appears quite promising, as it has been described as useful to elongate the duration of cell growth in nutrient restricted environments[17].

**Figure 6:**
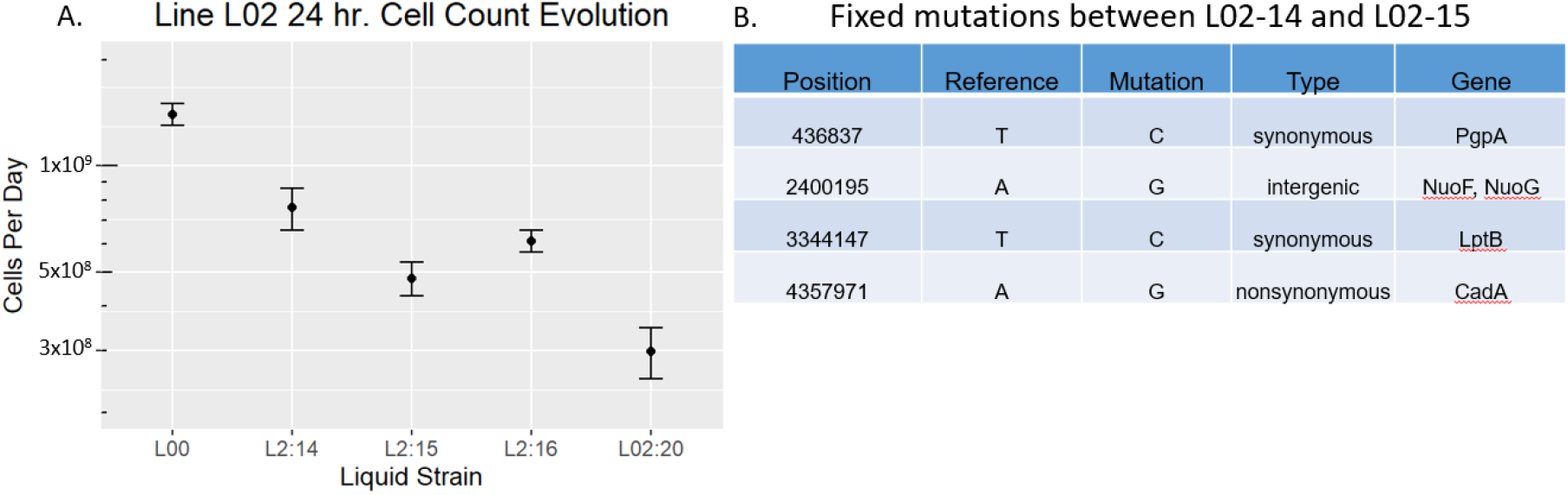
Decline in 24hr cell count of Liquid MA strain 2. A. 24hr CFU estimates of cell count in a time series of strain L02, with a focus on the drop between day 14 and day 15. B. Four mutations were fixed between day 14 and 15.

## 4 Discussion

In summary, the liquid MA largely recapitulates the findings of a plate MA for MMR-E coli, with several modest differences. This result supports the perspective that MA experiments, even those which have been run on agar plates, provide reasonable estimates of organism mutation rates. It would appear that *E. coli* is not significantly perturbed in mutation rate by growing either in liquid or on plate colonies, over the course of 24 hours. Despite some fraction of the cells being nutrient deprived for 6-14 hours[26], *E. coli* appears evolved to manage this environmental challenge with no significant change to its genome-wide mutation rate. Given the expectation that bacteria routinely encounter nutrient deprivation for hours, or perhaps even days or months in their evolutionary past, this result is not exceptional.

A primary finding of this research is that the liquid MA, single-cell bottlenecking by serial dilution does indeed work. By human hand, the task of streaking 16 samples, no more than 16 minute’s work even for the most un-familiar, turns into a task of about 90 minutes for a liquid MA, a poor incentive for anyone considering replicating the work presented in this article. However, the potential strength of the liquid MA will lie in it’s scalability, in the promise of automation, which could in principle require no more than picking which well a researcher would like to load per sample into a robot for serial dilution. In the hands of a robot, we expect that contamination would be somewhat less of a risk while running MA experiments, though we note that proper biosafety cabinet technique by hand also results in no risk of contamination. Given that some organisms do not grow well on agar plates, this protocol may also provide some researchers with an avenue to pursue questions of variation in mutation rate which have previously been entirely intractable.

We note also the overall resilience of the standard MA in the face of some criticisms to the technique over the years. Despite fear of induced selection or bias on behalf of the colony picker, our cursory examination of MA results between liquid and plate demonstrate the reliability of both protocols. We do not detect a large difference in mutation rate, or in mean fitness cost per mutation. One minor benefit of running a liquid MA is a less variant number of cell divisions per day per line. This may result in a more accurate estimation of mutation rates, though if true the effect is modest, if present at all: we detect a difference of 2% between the estimates of mutation rate between liquid and plate MA’s.

The statistically significant, though modest difference in mutation spectrum is curious, and for this we have no immediate explanation. Before putting much weight in speculation, we would like to see the result reproduced. With this preface, we note that the liquid mutation rate is skewed even more strongly to A:T→ G:C than the previous results of plate MA experiments (p = 8.8e-8, *χ*-square), to a ratio of roughly 4.7:1. The most parsimonious mechanism for this mutation spectrum of transitions is base-pair tautomerization, as proposed by Watson and Crick in 1953[27] and iterated upon since[21].

Watson and Crick noted that the temporary shift of hydrogen atoms to neighboring atoms might cause some level of nucleotide mispairing. They hypothesized that the A:T →G:C mutation ought to be more frequent than the G:C→ A:T, because the former requires a single hydrogen atom shift, while the latter requires two hydrogen atom shifts. In is unclear if or why the the medium of growth would alter the ratio of tautomerization, though it is known that the spent culture of *E. coli* significantly changes in pH (increase to pH of 9) and chemical composition[1, 19] which also affects gene expression profiles. Further, the decreased G:C→ A:T rate may be partly explained by the E coli of the liquid MA being exposed to less oxidative stress resulting in fewer damaged guanosine residues, a base prone to oxidative damage[10, 6]. However, growth of wild-type *E. coli* in the complete absence of oxygen mildly increased mutation rates, on the order of 2-fold, with no reduction in mutations to guanosines[20]. Regardless, the recurrent theme of MMR- *E. coli* and its derived strains are that the mutation spectrum is skewed in favor of transitions, as expected[12, 28]. The data of this work and prior MMR-growth experiments lend credence to the idea that tautomerization, if rare, is likely biologically relevant, such that life evolves DNA-repair pathways to manage its occurrence. We note that the degree of the imbalance in mutation types is potentially quite useful in the verification of novel sequencing technologies seeking to detect DNA mutations, to ensure that they are working properly.

With regard to fitness estimates of *E. coli*, discrepancies between the 96-well plate reader OD and actual cell count is a known issue[18], but with reiteration will hopefully dissuade researchers from relying on 96-well plate readers overmuch, in the context of mutation-accumulation experiments. As to why the discrepancy exists, at least in this experiment, we propose that the mutation-burdened cells of a MA experiment are more prone to biofilm formation or flocculating, sinking to the bottom of a well containing liquid growth medium. Either phenotype results in a layering of cells, which may lead to increased occlusion of the OD detector and yielding unusually high OD readings. Regardless of the ultimate cause of the outlier readings, the conclusion remains that attempting to quantify relative carrying capacity from OD600 in plate readers from cells with many accumulated mutations is at best done with caution, and should be backed up with CFU measurements.

